# Biphasic curvature-dependence of cell migration inside microcylinders: persistent randomness versus directionality

**DOI:** 10.1101/2022.12.30.522287

**Authors:** Xiaoyu Yu, Haiqin Wang, Fangfu Ye, Xiaochen Wang, Qihui Fan, Xu Xinpeng

## Abstract

Cell-scale curvature plays important roles in controlling cell and tissue behaviors. However, these roles have not been well quantified, and the underlying mechanisms remain elusive. We combine experiments with theory to study systematically the curvature-dependence of cell migration inside PDMS microcylinders. We find that persistence is positively correlated with speed, following the universal speed-persistence coupling relation, *i*.*e*., faster cells turn less. Cell migration inside microcylinders is anisotropic and depends on curvature in a biphasic manner. At small curvatures, as curvature increases, the average speed and anisotropy both increase, but surprisingly, the average persistence decreases. Whereas as the curvature increases over some threshold, cells detach from the surface, the average speed and anisotropy both decrease sharply but the average persistence increases. Moreover, interestingly, cells are found to leave paxillins along their trajectories (on curved but not planar surfaces), facilitating the assembly of focal adhesions of following cells. We propose a minimal model for the biphasic curvotaxis based on three mechanisms: the persistent random “noise”, the bending penalty of stress fibers, and the cell-surface adhesion. The findings provide a novel and general perspective on directed cell migration in the widely existing curved microenvironment of cells *in vivo*.

---

Geometrical cues at various length scales play important roles in guiding cell and tissue behaviors^1–4^. For example, curved topographical features at nano- and micro-scales have long been known to affect cell fate and differentiation^5–9^, cell migration^10–12^ and alignment^12–14^. Particularly, cell migration directed by anisotropic surfaces with subcellular-scale curvatures has been extensively studied and is widely known as “contact guidance”^15^. Recently, it has been understood that cell-scale curvature can also direct cell and tissue behaviors^16,17^, termed as “curvotaxis”^11^. However, the mechanisms underlying curvotaxis remains elusive^11^, particularly for cells migrating on cell-scale concave surfaces^10,14,16^. Concave surfaces are ubiquitous in many *in vivo* contexts. For example, during embryonic development, cells migrate from the hypoblast to line the blastocyst cavity, forming a primary yolk sac^18,19^. In neutrophil extravasation, neutrophils attach and crawl on the endothelial lumen surface, preparing for further transmigration^20^. In pancreatic ducts subject to oncogenic transformation, the direction of tumor growth depends on the radius of duct curvature: tumors expand outwards on narrow ducts (exophytic), while they grow inwards on larger ducts (endophytic)^21^. Moreover, human fibrosarcoma cells have also been observed *in vivo* to migrate inside the microvessels and capillaries^22^.

Over the last five years, many *in vitro* experiments have been carried out to unveil the mechanisms underlying cell curvotaxis on concave surfaces. For example, Bade *et al*^10^ observed the migration of fibroblasts on a sphere-with-skirt surface and found that cells form “chords” across the concave gap to avoid SF bending. Werner *et al*^12^ later studied the migration of human bone marrow cells (hBMSCs) on semi-cylinders and also observed the formation of SF “chord” structures in cells on concave surfaces. This mechanism was then used to explain the isotropic and weakly persistent migration of hBMSCs on concave surfaces, which is in strongly contrast to anisotropic persistent migration of hBMSC on the convex surface of the semi-cylinders. Moreover, some recent experiments find that cells on some complex curved surfaces try to avoid convex region and migrate directionally towards the concave region^11,12,23^.

On the other hand, to understand the mechanisms underlying curvotaxis, several hypothesis and theoretical models have been proposed. As early as in 1976, Dunn and Heath hypothesized that the orientation and migration of fibroblasts along the axis of cylindrical fibers (of micrometer radius) is because SFs cannot assemble or operate in the bent conditions (*i*.*e*., the bending penalty of SFs). Whereas this hypothesis does not explain the orientation of epithelial cells along lateral directions perpendicular to the cylinder axis^13,14,24^. To explain the different orientation of different cells on convex cylinder surfaces, Biton and Safran^25^ proposed a phenomenological model based on the competitions between the actomyosin contractility and the bending penalty of SFs and cell body. However, this model has not yet been used to study curvotaxis on concave surfaces and it can only predict static orientation of cells but not the dynamic cell migration. More recently, some simulations based on agent-based models have been done^23,26,27^. He and Jiang^26^ found that the constraints of curved substrates bias the direction of the stochastic protrusion force. Particularly, concave substrates facilitate protrusion force along the long axis of a cylindrically curved substrate and promote more persistent migration. Vassaux *et al*^23^ highlighted the importance of nucleus as a curvature sensor; the displacement of the nucleus drives directional cell migration toward concave regions. Despite capturing various experimental observations, these models have thus far primarily provided qualitative insights on cell curvotaxis and the underlying physical mechanisms for curvotaxis are still controversial. A quantitative statistical analysis on the curvotaxis is missing and is necessitated to evaluate the different mechanisms.

In this work, we combine *in vitro* experiments with theory to study systematically and quantitatively the curvotaxis of two distinct types of cells (fibroblasts, NIH3T3, and epithelial cells, MCF10A) on the inner concave surfaces of polydimethylsiloxane (PDMS) microcylinders (see Fig. 1). In comparison to complex curved surfaces used in most recent curvotaxis studies^10,11^, the major advantages of using cylinders to study curvotaxis are two-folded. Firstly, the cylinder is the simplest curved surface and it does not have intrinsic (Gaussian) curvature. Technically, cylinders are also easy to fabricate, and many periodic arrays of homogeneous PDMS cylinders have been fabricated. With a large volume of cell trajectories obtained using confocal microscopy and multi-cell tracking, we are able to provide a statistical description of cell curvotaxis. Secondly, our microcylinders are with closed tube-like geometry, which are different from open semi-cylindrical surfaces used in previous studies^12^. Such closed geometry better mimics natural concave surfaces such as glands or vessels existing *in vivo* and avoids artificial boundary effects, which may arise, for example, due to protein unfolding and receptor-ligand accessibility^28^.

**Fig. 1.**
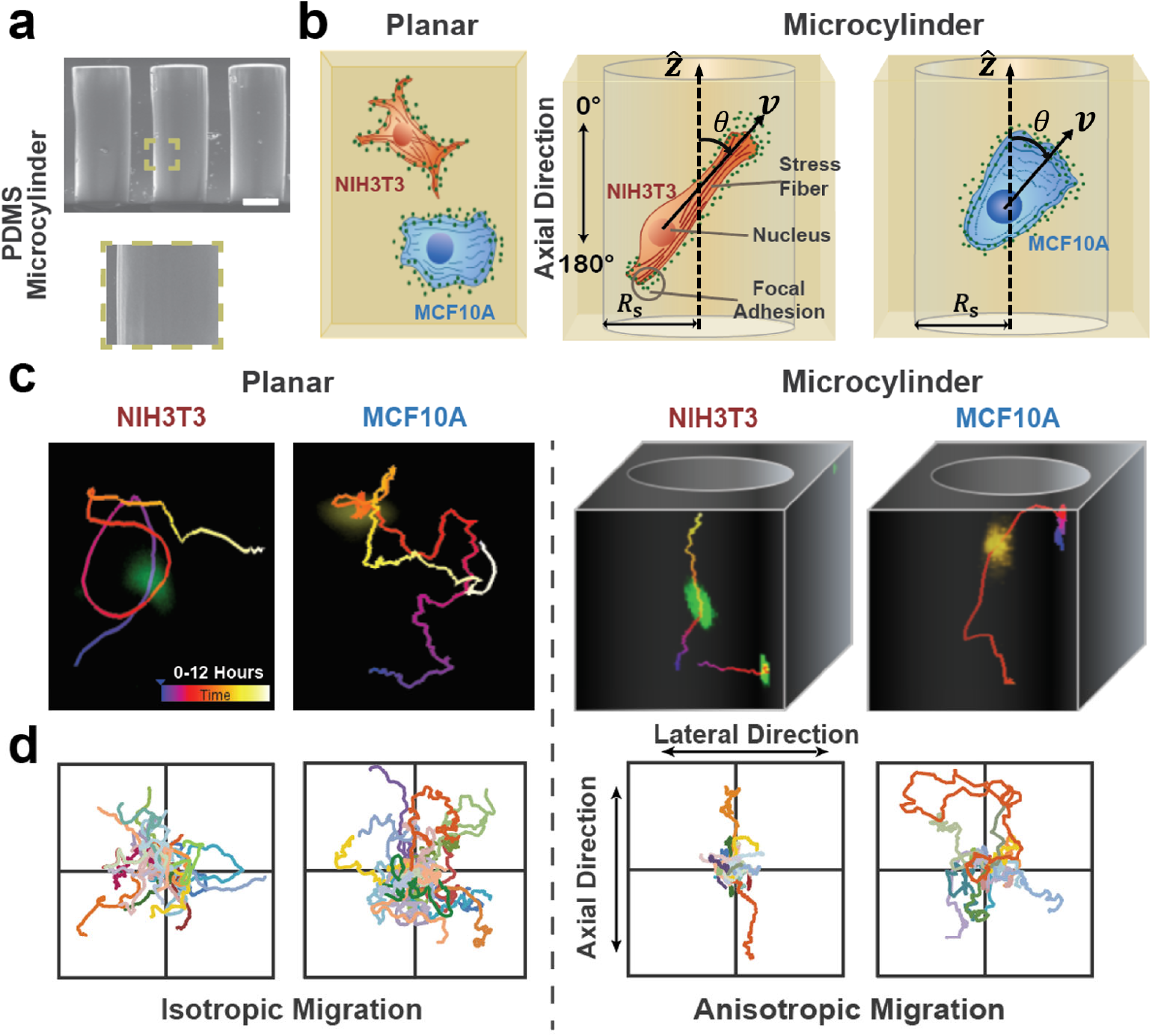
PDMS microcylinders and representative trajectories of cells migrating on planar surfaces and on the inner concave surfaces of microcylinders. (a) Scanning electron microscopy image of the stiff PDMS microcylinders. Scale bar: 100μm. The enlarged picture illustrates the micron-scale smoothness of the microcylinder surfaces. (b) Schematic diagram for the migration of fibroblasts and epithelial cells on planar surfaces and inside microcylinders (where the two principal curvatures are: 0 and a non-zero *κ*_*S*_ with curvature radius 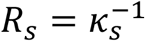). The orientation of cell migration inside microcylinders is described by the angle *θ*with respect to the axial 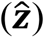 direction. (c, d) Representative trajectories of individual epithelial cells (left, MCF10A) and fibroblasts (right, NIH3T3) migrating on planar surfaces and inside microcylinders. The random migration of both types of cells is isotropic on planar surfaces but becomes highly anisotropic inside microcylinders and aligns averagely along the cylinder axial direction. The curvatures *κ*_*S*_ of microcylinder in (a) and (c) are both 1/150 μm.

We find that for both fibroblasts and epithelial cells, the concave cylinder surfaces bias the persistent random migration of the cell and directs cell migration along cylinder axis. The curvature-dependence of cell migration is biphasic and for curvature over some threshold, cells detach from the surface and SF chords are formed. Interestingly, cells on concave surfaces leave paxillins along their trajectories (but not on planar surfaces), which facilitates the assembly of focal adhesions of following cells. We propose a minimal theoretical model for the biphasic curvotaxis based on three mechanisms: the persistent random “noise”, the bending penalty of stress fibers, and the cell-surface adhesion. Our findings unveil the dominant mechanisms underlying curvotaxis on concave surfaces, provide a general perspective on the directed cell migration in the widely existing curved microenvironment of cells *in vivo*, and give insights into the geometric design of artificial scaffolds for tissue engineering and regenerative medicine.

## Characterization of cell speed and persistence on surfaces

We track and analyze the 3D migration trajectories of fibroblasts (NIH3T3), and epithelial cells (MCF10A-GFP) on the planar surfaces and on the inner concave surfaces of polydimethylsiloxane (PDMS) microcylinders (as shown schematically in Fig.1b). The two representative types of cells have distinct SF organization. Fibroblasts when polarized usually have pronounced thick bundles of straight SFs and have been found to orient longitudinally on convex cylinders^13^. In contrast, epithelial cells have relatively thin SFs, and, therefore, are more difficult to be visualized (as shown schematically in Fig. 1b and in Fig. 4a) and known to align laterally perpendicular to the axis on convex cylinders^13^. However, when these cells migrate on concave surfaces, we will discuss later that their distinct SF organization make no qualitative but only quantitative difference in their curvotactic migration behaviors.

The PDMS microcylinders are fabricated using the soft-lithography technique (Fig.S1a) and have nano-level smooth inner surfaces (see Fig.1a), allowing for high-throughput observations (Fig.S1b). The PDMS microcylinders are stiff (with Young’s modulus about 750 kPa) and the two anisotropic principal curvatures are: 0 (along the axial direction 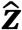) and *κ*_*S*_ (along the lateral direction with *κ*_*S*_ < 0 for inner concave surfaces and the curvature radius denoted by 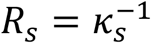), respectively, see Fig. 1b. In our experiments, periodic arrays of homogeneous PDMS cylinders have been fabricated with each array having a height about 400 μm, providing sufficient space for migration of individual cells, and an average curvature radius *R*_*S*_, ranging from 50 μm to 200 μm.

The cell migration on planar surfaces and inside microcylinders are tracked and recorded for about 12 hours by confocal laser scanning microscope equipped with an on-stage incubator. Based on the recorded cell trajectories, we analyzed three major quantities: the instantaneous migration speed, *v*, the cell migration orientation, *θ*(defined relative to the cylinder axis, as shown in Fig. 1b), and the persistent time, *τ*_*p*_. Technically, the instantaneous velocity ***v*** (including *v* and *θ*) is obtained from the displacements of two successive frames (with 4 minutes intervals) in the trajectories (see Figs. 1c, d). The persistent time, *τ*_*p*_, is calculated as the time duration for the cell to change its migration direction by a turning angle about 90^°^ (see Supplementary Information). Both fibroblasts and epithelial cells migrate on planar surfaces in a random and isotropic manner, following a persistent random walk with *v*∼0.5 μm/min and *τ*_*p*_∼30 mins (See Fig. S10 in the Supplementary Information). In contrast, although cylinders do not have intrinsic (Gaussian) curvature, cell migration inside microcylinders shows a strong and complex dependence on the curvature, *κ*_*S*_, of the cylinder.

For the two types of cells migrating inside microcylinders with a given curvature *κ*_*S*_ (ranging from 0 to 1/50 μm^-1^), we firstly find that both *v* and *τ*_*p*_ follow exponential probability distribution (Figs. 2a, c). The exponential speed distribution has been claimed to represent a universal feature of cell migration on solid surfaces^29,30^. In contrast, the exponential probability distribution of persistent time, *τ*_*p*_, has rarely been demonstrated. Note that such exponential distribution of *τ*_*p*_ is usually assumed in the run-and-tumble model of swimming bacteria to produce a constant tumbling rate". Secondly, for cell migration on the surface with a given curvature *κ*_*S*_, we find that the mean instantaneous speed *v* and the instantaneous persistence time *τ*_*p*_ are exponentially positively correlated, following the universal speed-persistence coupling (UCSP) relation^32^: *τ*_*p*_ ∝ *e*^*λv*^ (see Fig. 2e) with *λ* being a fitting correlation parameter. That is, when cells migrate fast on concave surfaces with a given curvature, they usually have high persistence and turn less frequently. The UCSP relation is firstly identified for the migration of cells adhered on 1D and 2D planar surfaces, as well as for the migration of cells embedded within 3D microenvironment^32^. Such exponential positive correlation has been suggested to result from the coupling of retrograde flow to the asymmetry of polarity cue molecules^32^. Here for cells migrating on concave surfaces, we find that the fitted correlation parameter *λ* depends on the curvature *κ*_*S*_ in a biphasic manner (see Fig. 2f). At small curvature, *λ* decreases with *κ*_*S*_ but as curvature is larger than some threshold, *λ* increases again.

**Fig. 2.**
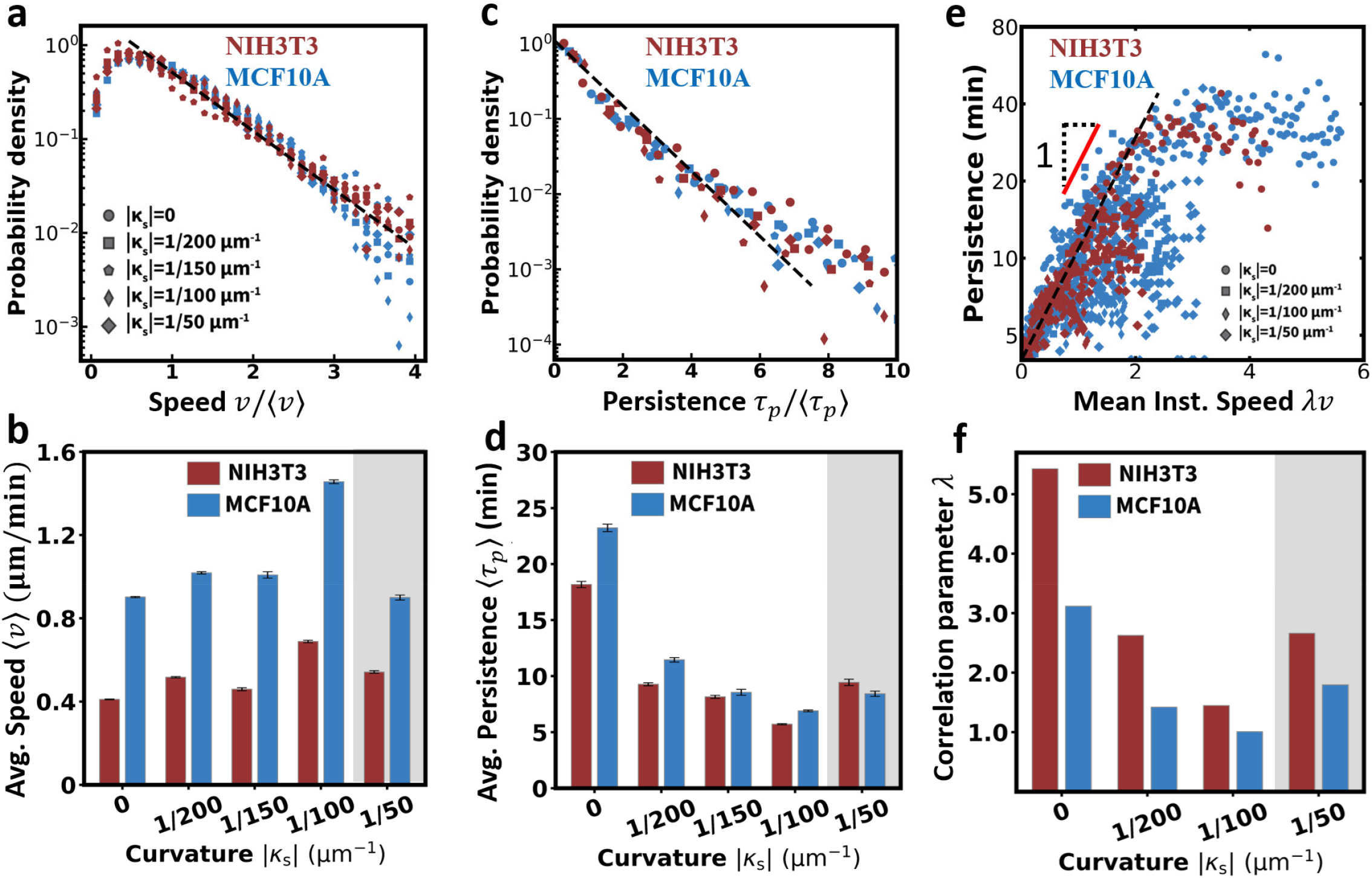
Cell speed and persistence (distributions and correlations) for cells migrating on surfaces and their biphasic dependence on surface curvature, *κ*_*S*_ (particularly, *κ*_*S*_ = 0 for planar surfaces). (a) The universal exponential probability density function (PDF) of instantaneous speed *v* (normalized by its average ⟨*v*⟩) for various *κ*_*S*_. (b) The average speed ⟨*v*⟩ is plotted as a function of *κ*_*S*_. (c) The universal exponential PDF of persistence time *τ*_*p*_ (normalized by its average ⟨*τ*_*p*_⟩) for various *κ*_*S*_. (d) The average persistence time *τ*_*p*_ is plotted as a function of *κ*_*S*_. (e) The universal speed-persistence coupling (UCSP) relation. (f) The curvature-dependence of the fitted UCSP correlation parameter *λ*. Here two types of cells are studied: epithelial cells (MCF10A, blue) and fibroblasts (NIH3T3, red).

Surprisingly, however, we find that cells inside microcylinders have larger average speeds, ⟨*v*⟩ (*i*.*e*., migrate faster) but smaller average persistent time, ⟨*τ*_*p*_⟩ (*i*.*e*., turn more frequently) in comparison to their migration on planar surfaces (see Figs. 2b,d). This negative correlation between ⟨*v*⟩ and ⟨*τ*_*p*_⟩, seemingly contradictory to the UCSP relation between *v* and *τ*_*p*_, is actually consistent with the biphasic dependence of *λ* on *κ*_*S*_ and by the inverse proportionality of *λ* with ⟨*v*⟩ (see Fig. S4 in the Supplementary Information). The parameter *λ* is smaller for cell migrations inside microcylinders than that on planar surfaces, and hence although ⟨*v*⟩ is larger yet ⟨*τ*_*p*_⟩ can still be smaller. In this work, instead of resolving the underlying complex biochemical mechanisms and explaining the curvature-dependence of cell speed and persistence, we focus on the migration anisotropy and present a coarse-grained phenomenological description to the curvotaxis of cells migrating inside microcylinders by considering the cell as a self-propelled soft-matter object that is constrained geometrically to the curved cylinder surfaces.

### Anisotropically persistent random cell migration

From the cell migration trajectories, we have also examined the dependence of cell orientation on the microcylinder curvature, *κ*_*S*_. As shown in Fig. 3a, we see that for both fibroblasts and epithelial cells, the distribution of cell migration orientation on planar surfaces (with *κ*_*S*_ = 0) is highly isotropic. In contrast, cell migration on the inner concave surfaces of microcylinders becomes anisotropic, aligning along the axial direction. Such anisotropy in cells’ migrating orientation can be quantified by the nematic order parameter defined via

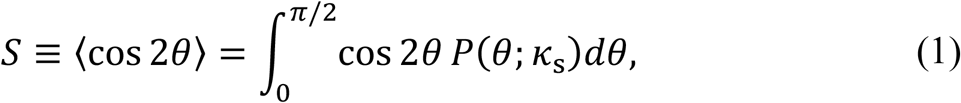

in which ⟨… ⟩ represents the statistical average, and *P*(*σ; κ*_s_) is the probability distribution of the orientation *θ*of cells migration inside the microcylinder with curvature *κ*_s_. In practice, the statistical analysis is carried out using both time and ensemble averages over the cell trajectories. In Fig. 3b, the nematic order parameter *S* are calculated for the two types of cells migrating on surfaces of different curvatures. From Figs. 3a-b, we find that the dependence of cell migration anisotropy on microcylinder curvature *κ*_*S*_ is biphasic: at small curvature, the orientation anisotropy characterized by *S* increases with *κ*_*S*_, while for curvature larger than some threshold, |*κ*_th_| (about 1/50 μm^-1^, the orientation anisotropy decreases sharply and the cell migration switches to be more isotropic again.

**Fig. 3.**
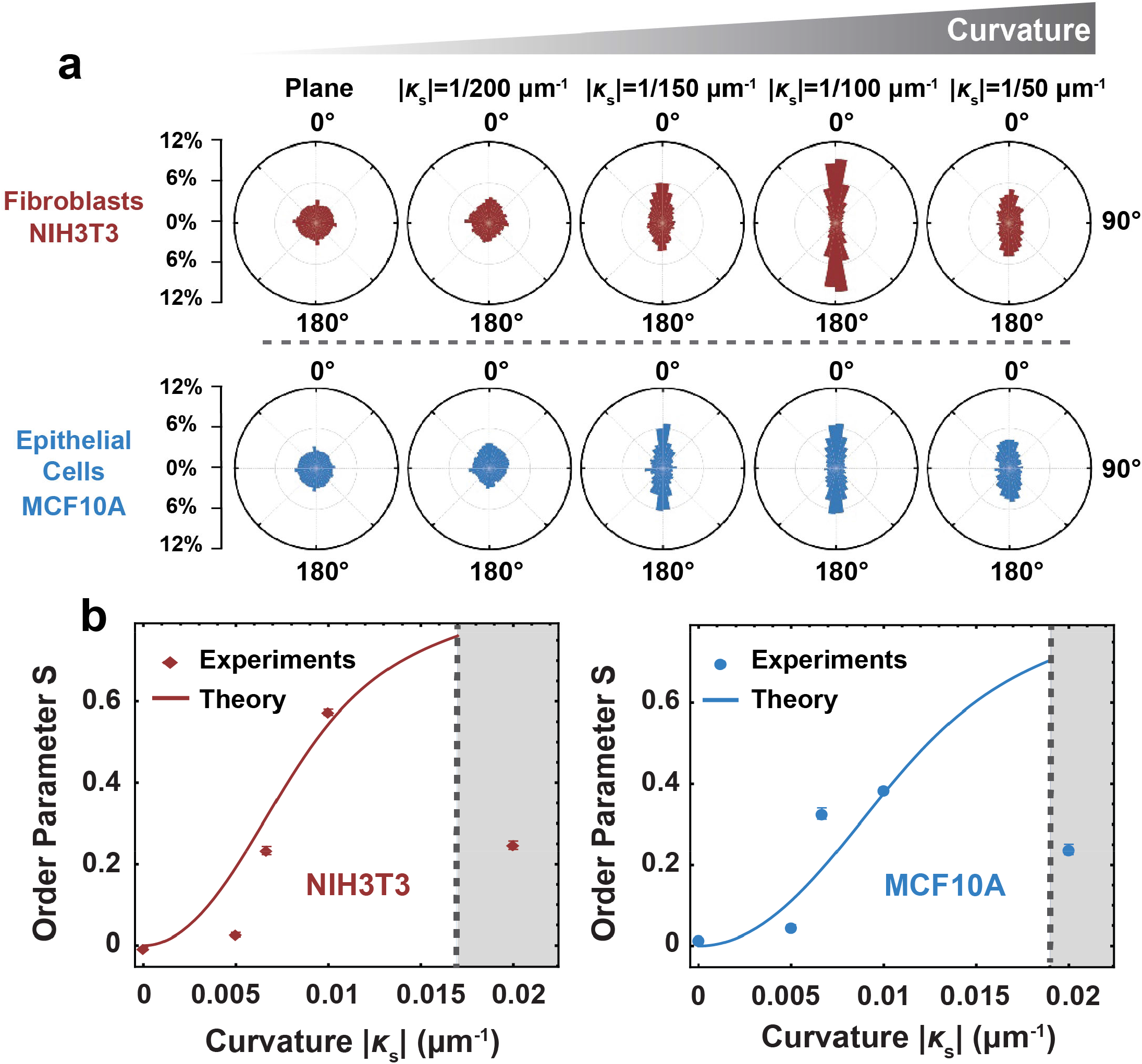
Curvature-directed anisotropic cell migration on surfaces of various curvature, *κ*_*S*_ (particularly, *κ*_*S*_ = 0 for planar surfaces). (a) Probability distribution of migration orientation *θ* for various *κ*_*S*_. (b) The migration anisotropy is quantified (dots) using the nematic order parameter *S* = ⟨cos 2*θ*⟩ defined in Eq. (1). The theory (solid line) fits the experiments well using one fitting parameter ℬ that characterizes the different bending stiffnesses of the F-actin stress fibers for different cell types relative to the persistent random cell migration. Here ℬ is fitted to be, 30 and 41, for epithelial cells (MCF10A, b) and fibroblasts (NIH3T3, c), respectively.

### Curvature-dependent organization of SFs, nuclei, and FAs

To unveil the mechanisms underlying the biphasic curvotaxis inside microcylinders, particularly how surface curvatures control the speed, persistence, and orientation of cell migration, we image and analyze the organization and distribution of F-actin SFs, cell nuclei, and focal adhesions (FAs). Firstly, we consider cells migrating on microcylinders with small curvature, |*κ*_s_| < 1/50 μm^-1^. For fibroblasts, it can be clearly seen (see Figs. 4a, b) that the F-actin SFs are randomly distributed in cells on planar surfaces, while the SFs tend to align strongly along the cylindrical axis in cells inside microcylinders. Moreover, from the side view shown in Fig. 5a, fibroblasts adhere tightly to the concave surfaces and hence the SFs are bent accordingly. Therefore, SFs reorient and align naturally to the less bent direction close to the axial direction where the bending penalty of SFs is minimized. In contrast, the SFs of epithelial cells are much thinner and diffusive (see Fig. 4a). However, it is still clear that the SFs of epithelial cells inside microcylinders become much thicker (easier to visualize) than those on planar surfaces and are aligned to the axial direction to avoid their bending. The distinct SF structure of the two types of cells underlie the quantitative differences in their motility as shown in Fig. 2 and Fig. 3 such as speed, persistence, and the degree of curvature-induced alignment. The curvature-induced alignment of SFs could promote actin turnover and improve the efficiency of intracellular molecule transport^33,34^.

**Fig. 4.**
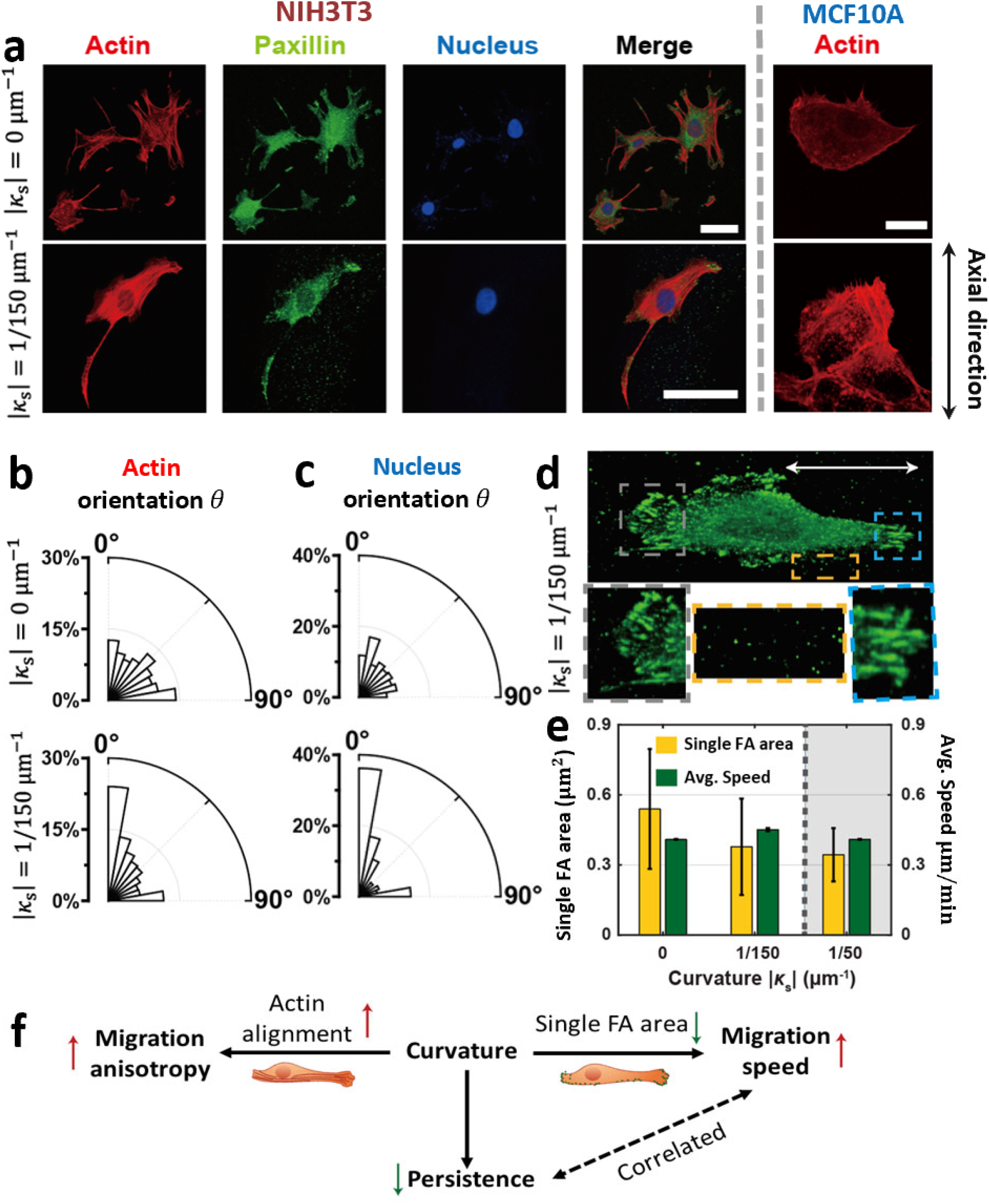
Cell-scale curvature directs cell migration by inducing the reorganization of the F-actin stress fibers (SFs), cell nucleus, and focal adhesions (FAs). (a) Fluorescent images of F-actin, paxillin, and nucleus for fibroblasts (NIH3T3, the first four columns) and images of F-actin for epithelial cells (MCF10A, the last column) on planar surfaces (top) and inside microcylinders (bottom, |κ_*S*_| = 1/150μm^-1^), respectively. (b-c) The probability distributions of the orientation of (b) F-actin and (c) nucleus in fibroblasts on planar surfaces and inside microcylinders. (d) The expression of paxillin in and out of the fibroblast inside a microcylinder of |κ_*S*_| = 1/150μm^-1^. (e) The area of individual FAs and the average speed ⟨*v*⟩ of fibroblasts migrating on surfaces of various curvature κ_*S*_. (f) Graphical summary of the physical mechanisms underlying the curvature-directed cell migration and alignment inside microcylinders.

**Fig. 5.**
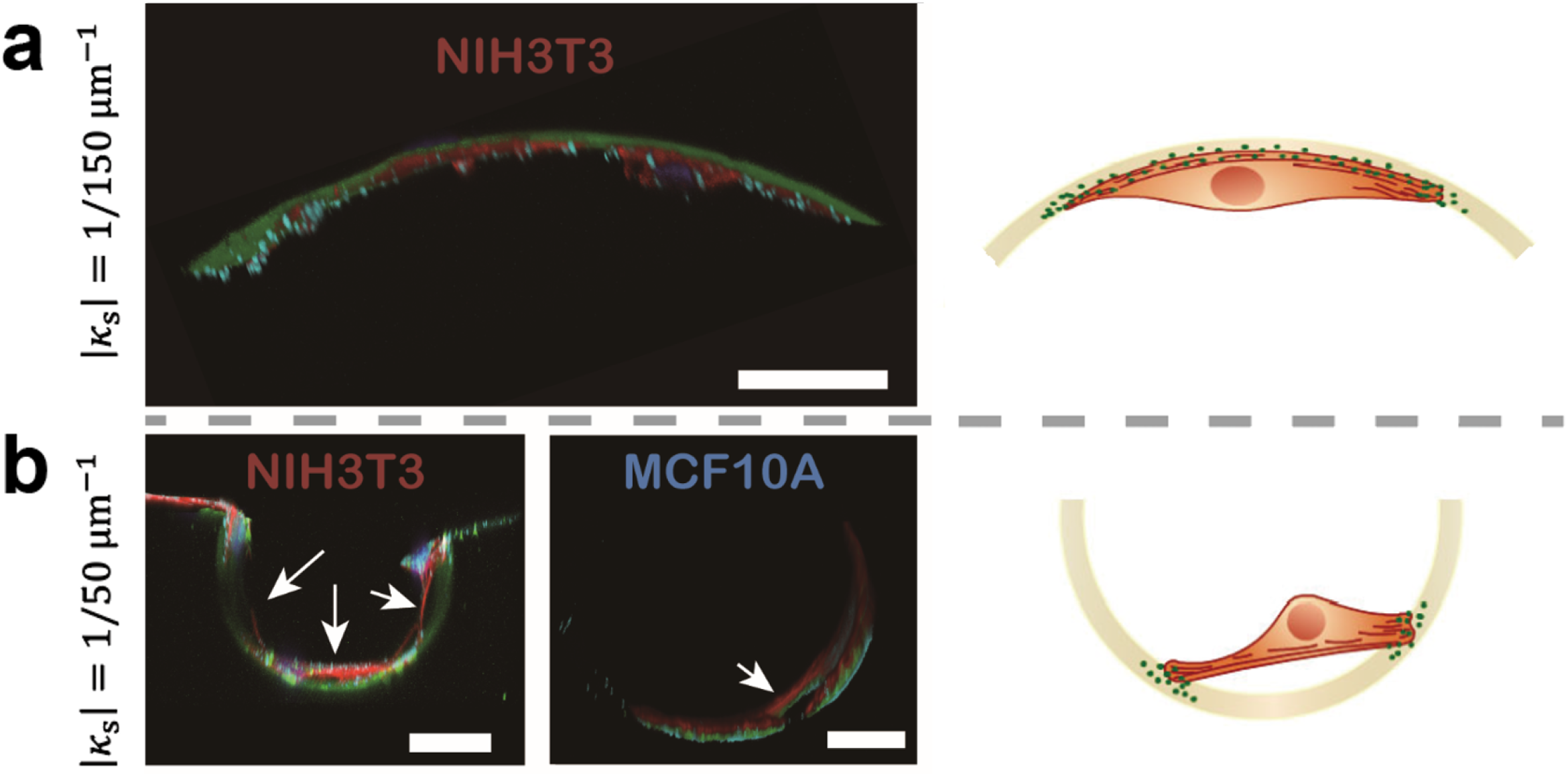
Detachment of cells and chord formation of apical SF bundles inside the microcylinder when its curvature |κ_*S*_| is larger than some threshold around 1/100 μm^-1^. **(Left panel)** Side view of the fluorescence images of F-actin (red), focal adhesion (blue), and surface (green) for cells inside microcylinders with (a) |κ_*S*_| = 1/150 μm^-1^ (adhered state) and (b) |κ_*S*_| = 1/50 μm^-1^(chord state), respectively. (Scale bars, 20 μm). **(Right panel)** Schematic representation of cell shape at the adhered state (top) and chord state (bottom), respectively.

Furthermore, the orientation of cell nuclei also changes with the underlying surface curvature (see Fig. 4c). The nucleus is round in cells migrating on planar surfaces but is elongated along the axial direction on concave cylinder surfaces. This implicates that the surface topography not only regulates cell migration but also directly impacts nucleus shape and orientation, which is known to regulate gene expression^11,23^ by controlling the nucleopore width and the chromatin condensation.

In addition, the structure changes of FAs are another important component for regulating cell migration. We characterize the distribution of FAs by labeling and imaging the paxillin. For fibroblasts on planar surfaces, paxillins are distributed randomly and isotropically (Fig. 4a). In contrast, for fibroblasts on concave surfaces (Fig. 4d), most paxillins are distributed densely around the cell protrusions and aligned in the axial direction with few small paxillins distributed sparsely in lateral directions. Moreover, the sizes (or area) of single FAs are also found to be ∼1/3 smaller in cells adhered on cylinder surfaces than those in cells on planar surfaces (Fig. 4e). We can then conclude that the curvature-induced increase in molecule transport due to SF alignment and the decrease in FAs sizes act together to promote cell migration and increase the average speed ⟨*v*⟩.

Interestingly, we note that on concave surfaces, paxillins are left over and scattered onto the cylinder surfaces outside the cell along its migration trajectories (Fig. 4d). Such curvature-induced unusual distribution of paxillins may explain some unique motility behaviors of cell curvotaxis. For example, the paxillins left on curved surfaces may assist the following cells to form new FAs easier and hence change migration directions easier, which then explains the decrease of the fitted UCSP correlation parameter *λ* and the average persistence time ⟨*τ*_*p*_⟩ on curved surfaces.

Secondly, when the cylinder curvature is large and exceeds some threshold (for example, |*κ*_th_| 1/50 μm^-1^), both types of cells are found to detach partially from the concave cylinder surfaces and the apical SFs form a “chord” structure (Fig. 5b). Accordingly, the anisotropy of cell migration has a sharp decrease (Fig. 3). This further addresses the essential roles of surface curvature in controlling the organization of SFs and regulating cell migration orientation.

### A minimal model: persistently randomness vs. directionality

The migration of cells on planar surfaces has long been described by persistent random walk (PRW)^35,36^. Here we have also confirmed that the migration of both fibroblasts and epithelial cells on planar PDMS surfaces can be well described by the following alternative form of PRW model^37^: 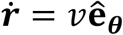and 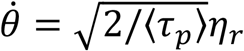, in which the superscript dot denotes the time derivative, *v* is the instantaneous cell speed, **ê**_0_ = (cos *θ*, sin *θ*) is the unit vector of instantaneous cell velocity on the surfaces with *θ*being the angle relative to some arbitrarily chosen direction, *τ*_*p*_ is the persistence time with ⟨*τ*_*p*_⟩ being its average, and *η*_*r*_ represents the white noise with zero mean and unit variance. Note that here *v* follows an exponential distribution with an average ⟨*v*⟩, and *v* is exponentially positively correlated with *τ*_*p*_. Such speed-persistence coupling has recently been found to enhance search efficiency of migrating cells^38^. Using this correlated PRW model, we fit the mean-square-displacements and velocity-autocorrelation-functions calculated from cell trajectories and obtain that for fibroblasts (NIH3T3), ⟨*v*⟩ 0.41 μm/min and ⟨*τ*_*p*_⟩ 32.9 min; for epithelial cells (MCF10A), ⟨*v*⟩ 0.90 μm/min and ⟨*τ*_*p*_⟩ 25.0 min.

For cells migrating inside microcylinders, the above correlated PRW model can be further modified to describe the curvature-directed anisotropic migration. Given the translational symmetry of microcylinders, we propose that the effects of surface curvature on the cell migration can be separated into two parts: the curvature-dependent migration speed *v*(*κ*_*s*_) and persistence time *τ*_*p*_(*κ*_*s*_); the curvature-induced torque that can be derived from a curvature-dependent potential, *U*(*θ*; *κ*_*S*_). In this modified PRW model, cell curvotaxis is described by the following over-damped Langevin equations:

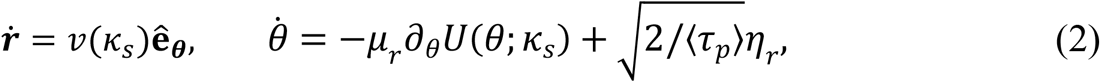

in which **ê**_0_ = (cos *θ*, sin *θ*) is the unit velocity vector with *θ*being the angle relative to the cylinder axial direction 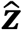(see Fig. 1b), and *μ*_*r*_ is the rotational mobility. Here *v* also follows the exponential distribution and is exponentially positively correlated with *τ*_*p*_, but the average ⟨*v*⟩ is a larger constant and ⟨*τ*_*p*_⟩ is a smaller constant than those of cells on planar surfaces. Note that generally the curvature-dependent potential should also depend on the position, *i*.*e*., *U*(***r***, *θ*; *κ*_*S*_), for cell curvotaxis on surfaces of more complex topography. This modified correlated PRW model provides a generic phenomenological description of cell migration, in which the competition between persistent randomness and directionality induced by asymmetric cues is a ubiquitous and important component.

To find *U*(*θ*; *κ*_*S*_), we propose a phenomenological model of cell as a thin elastic circular disk that is reinforced by aligned SFs and adhered to the inner constraining (concave) surfaces of microcylinders. Following the classical plate theory^39^, we obtain the total bending energy of the adhesive cellular disk as (see Supplementary Information)

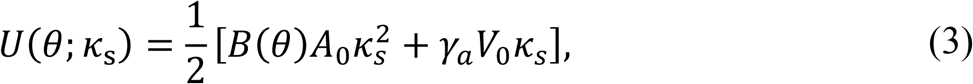

with the curvature *κ*_*S*_ < 0 for concave surfaces, the effective bending rigidity *B*(*θ*) = *B*_b_ + *B*_f_*L*_f_ sin^4^ *θ* /*A*_0_ depending on the orientation angle, *θ*, of SFs. Here 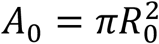, *V*_0_ = *A*_0_*h*_0_, *h*_0_, and *R*_0_ are the area, volume, thickness, and radius of the elastic cell disk, respectively. *B*_f_ and *L*_f_ are the bending modulus and total length of SFs, respectively. *B*_b_ and *γ*_*a*_ are the bending modulus and the adhesion energy density (per unit area) of the cell disk on the microcylinders. The adhesion energy density *γ*_*a*_ is positive, representing the preference of the adherent cells to spread out, which is analogous to the spreading coefficient of droplets on a complete wetting surface. Note that one should distinguish adhesion energy from interfacial energy as firstly clarified by P. G. de Gennes^40^; the adhesion energy is used here to predict the threshold curvature for cell detachment from underlying concave surfaces as discussed in the end of this subsection. In addition, we have assumed that the direction of cell migration inside microcylinders and planar surfaces coincides with the orientation direction of apical SFs. This assumption is not true in general, and the two directions have been found to deviate from each other significantly in a more complex sphere-with-skirt surface near the negative Gaussian curvature region^10^.

From *U*(*θ*; *κ*_s_), we can see that the average orientation of SFs and the cell migration are determined by the minimization of bending penalty of SFs as suggested in previous experiments^15,17^. Minimizing *U*(*θ*; *κ*_s_) gives *θ*= 0, or *π*, that is, the cells tend to align along the axial direction (with zero curvature) to minimize the bending penalty of SFs. Moreover, from Eq. (2) the distribution of cells’ migrating orientation is determined by the minimization of SFs bending penalty and the stochastic turning or tumbling of the cell migration. Particularly, the steady-state orientation distribution is given by a Boltzmann factor: *P*(*θ*; *κ*_*s*_) ∝ exp [*U*(*θ*; *κ*_s_)/*k*_*B*_*T*_*a*_], in which an effective temperature has been introduced via *k*_*B*_*T*_*a*_ ≡ 1/*μ*_*r*_⟨*τ*_*p*_⟩ to represent the strength of stochastic turning noise (in analogy to the temperature for the thermal noise in passive Brownian motion) with *k*_*B*_ being the Boltzmann constant. Furthermore, using this orientation distribution function and Eq. (1), we can calculate the nematic order parameter *S* to characterize the curvature-dependent anisotropic migration of fibroblasts and epithelial cells. Note that here we only have one fitting parameter 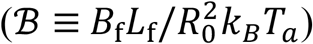that characterizes the strength of cells ’ preference to avoid the bending of SFs: for cells migrating on microcylinders with a given curvature, the larger ℬ of the cell is, the stronger its migration anisotropy is.

We first estimate the magnitude of ℬ using experimental values of relevant parameters in the literature. We assume their bending modulus *B*_f_ of SFs or F-actin bundles is proportional to the fourth power of the bundle diameter as that of elastic rods^39^. We use the bending modulus^25,41^ (∼10 μm · *k*_*B*_*T*) of a single F-actin filament (with a diameter of ∼8 nm) with *T* being the thermal temperature, and can then get an estimate of the bending modulus of F-actin bundles (of typical diameter ∼ 0.1μm) in cells to be *B*_f_∼10^5^ μm · *k*_*B*_*T*. Moreover, we assume the volume density of the aligned SFs to be proportional to the active contraction^25^ *ϵ*_*C*_, that is, the total length of SFs is given by *L*_f_ = *αϵ*_*C*_*V*_0_, in which *ϵ*_*C*_∼0.1, and *α*∼1 μm^-2^ is a positive proportionality constant with the dimension of inverse of length square. Using *h*_0_ = 0.1*R*_0_ and *R*_0_∼10 μm, we obtain *L*_f_∼10 μm, and assuming active temperature to be *k*_*B*_*T*_*a*_∼10^3^ *k*_*B*_*T*, we obtain 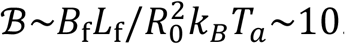. On the other hand, for the cell orientation characterized by the nematic order parameter *S* (Fig. 3b), we use our theoretical model to fit the experimental measurements and find ℬ to lie in the same order of 10. Particularly, we find that fibroblasts (with ℬ = 41) have stronger preference to avoid the bending SFs than that of epithelial cells (with ℬ = 30), which are consistent with the different SF structure of the two types of cells as discussed above.

When the cylinder curvature increases over some threshold, |*κ*_th_|, cells can detach partially from the cylinder by forming SF chords (see Fig. 5b, *i.e*., “chord state”). With the chord formation, the anisotropy of cell migration drops significantly and becomes almost isotropic (Fig. 3). Structure transition of SFs from bent state when they tightly adhered to the curving surface to the detached chord (unbent) state can be understood by our adhesive elastic-disk cell model. As cylinder curvature increases, the bending penalty of cell body and SFs become dominant over the energy decreased by adhering to the concave surfaces. In this sense, once the cylinder curvature is large enough, SF chords can form, which support cells back to morphology as they are on planar surfaces, resulting in a dramatic drop in migration anisotropy. However, if the principal cylinder curvature *κ*_s_ is only a little bit larger than *κ*_th_, the cells can detach and form chords only when cells’ orientation is very close to the circumferential direction where the cylinder curvature is larger than *κ*_th_. Moreover, even when |*κ*_s_| ≫ |*κ*_th_|, such non-uniform orientation-preference in the chord formation still exists around the axial direction. This mechanism, therefore, explains why the anisotropy of cell migration on microcylinders drops when |*κ*_s_| > |*κ*_th_| but is still non-zero (Fig. 3). Interestingly, we note that such an energetic-driven change of cell morphology on concave surfaces is similar to the Cassie-Wenzel transition of droplets on rough surfaces^42^. However, one should recognize that the underlying physical mechanisms are very different in the two seemingly similar phenomena. Adherent cells behave like an elastic solid while droplets only have interfacial energy, and the cell detachment from cylinder surfaces is determined by adhesion energy instead of interfacial energy as in droplets.

In addition, from our phenomenological cell model, we can also calculate the threshold curvature *κ*_th_ in terms of measurable quantities. Note that cells detach and form chords when the total energy of adhered cells on microcylinders, given in Eq. (3), becomes larger than the energy of the cell detached from the surface, given by *γ*_*a*_*A*_0_, which is approximated to be the energy needed to detach the circular-disk cell of area *A*_0_from planar surfaces. Based on this understanding, we find 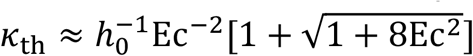, in which Ec = *ℓ*_ec_*/h*_0_ is the elastocapillary number and 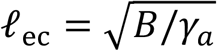is the elastocapillary length that characterizes the relative importance of bending energy of the cell and the adhesion energy. We can use the formula to estimate the order of magnitude of *κ*_th_ using experimental parameter values in the literature. We assume that the major contribution of bending modulus arises from the elastic bending of the cell body that scales as 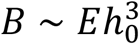, with *E* ∼ 10 kPa being the elastic modulus of the cell body. The adhesion energy density^41^ *γ*_*a*_ of cells is in the order of magnitude of *γ*_*a*_ ∼ 1 Pa · *μ*m. Using *h*_0_ = 0.1 *R*_0_ and *R*_0_ ∼ 10 *μ*m, we then obtain *ℓ*_e*c*_ ∼ 10^2^*μ*m and hence *κ*_th_ ∼ 0.01 *μ*m^-1^, which is in a good agreement with the threshold curvature observed in Fig. 3 and Fig. 5.

### Implications to curvotaxis on more complex curved surfaces

In this work, our experiments are carried out only for cell migration inside microcylinders. As explained by R. Feynman^43^, a bug that is living on a cylinder surface and only makes local measurements cannot discover that the surface is curved. However, here we find that cells can sense and respond to the cylinder curvature at the cell scale. Moreover, we would also like to point out that our minimal theoretical model and the proposed physical mechanisms are generic and can be easily extended to explain the rich curvotaxis behaviors observed in some recent literature^10,11^. For example, mesenchymal stem cells (MSCs) and fibroblasts on smoothly double sinusoidal surface^11,23^ are found to migrate towards the concave valleys and avoid the convex hills. In our model, the phenomenon is due to the competition of the SF bending penalty (by forming chords on concave surfaces) and the cell capability of having a larger contact area on concave surfaces due to the linear dependence of cell adhesion energy on the curvature as shown in Eq. (3). In another system, when fibroblasts migrate on sphere-with-skirt surfaces^10^, cells prefer to minimize SFs bending to avoid convex sphere cap (with positive Gaussian curvature) due to SFs bending penalty. The chord of apical SFs forms not around the inflection line (where the radial curvature is zero) but in a distance away from the line (where the radial curvature becomes a finite negative value); this can be explained by noting that SF chords form only when the curvature of concave surfaces is larger (in magnitude) than some threshold curvature *κ*_th_. After SF chords form on the skirt in the radial concave direction, fibroblasts are found to repolarize and migrate in the azimuthal direction. This can be easily understood in our phenomenological model: due to SF bending penalty, any forward or backward cell motion in radial directions will induce the SF transition from chord state to adhered state, causing larger resistance, and thus change the migration into azimuthal direction.

Our findings and the generic minimal model of single-cell curvotaxis could also provide insights into the orientation and migration of cells in aggregates such as confluent cell monolayers on curved surfaces^44–47^. The competitions among cell-cell junction, cortical actin bending, and cell contractility could result in even richer emergent collective behaviors^44^. Particularly, based on our single-cell models, for confluent cell monolayers on microcylinders, the cell-cell interactions may be modeled as nematic interactions in analogous to those in liquid crystals^48^. In this case, the cylinder curvature may provide essential polarized cues for cells and induce isotropic-to-nematic transition and rich defect dynamics. Therefore, this work paves the way to a quantitative understanding of curvotaxis at both single-cell and multicellular scales and may be beneficial for the geometrical design of artificial scaffolds in tissue engineering and regenerative medicine.

## Methods

### Fabrication of periodic arrays of PDMS microcylinders

The mask pattern is designed by AutoCAD software and processed into a chrome mask. Through the MA6 Mask Aligner (Karl Süss, Germany), the SU8-2150 (Microchem, USA) photoresist is exposed to templates. Drop 6μL of trichloro(1H,1H,2H,2H-perfluorooctyl) silane (Sigma) on a hot plate at 125 °C and place the template in a sealed environment formed by gas for five minutes to perform surface silanization. They were then held in place with a heat-curable silicone elastomer PDMS (Sylgard 184; Dow Corning). The entire structure was cured at 60 °C for 4 hours (stiffness Young’s modulus ∼750 kPa), cleaned with ethanol and dry nitrogen, oxidized inside an air plasma cleaner (Harrick Plasma), and used as is or coated with fibronectin (Sigma) at 50μg/mL in DPBS.

### Cell culture

NIH3T3 fibroblasts were cultured in DMEM (Corning) containing 10% BCS (GIBCO) and 1% penicillin-streptomycin (100X solution, Invitrogen No.15070-063) at 37°C in an atmosphere containing 5% CO_2_.

MCF10A non-tumorigenic epithelial cell line were cultured in DMEM/F12 (Invitrogen No.11039-021), Horse serum (Invitrogen No.16050-122), Pen/Strep (100X solution, Invitrogen No.15070-063), EGF (Peprotech,1 mg): Resuspend at 100 ug/ml in sterile ddH2O and store aliquots at -20°C, Hydrocortisone:(Sigma No.H-0888,1-g bottles): Resuspend at 1 mg/ml in 200-proof ethanol and store aliquots at -20°C, Cholera toxin:(Sigma No.C-8052,2-mg vials) resuspend at 1 mg/ml in sterile ddH_2_O and store aliquots at 4°C. Insulin (Sigma No. I-1882,100-mg vials): Resuspend at 10 mg/ml in sterile ddH_2_O containing 1% glacial at 37°C in an atmosphere containing 5% CO_2_.

### Immunofluorescence staining

For quantification of cell distribution. The cells were fixed with 4% paraformaldehyde for 10 minutes, permeabilized with 0.5% Triton™ X-100 for 10 minutes and blocked with 1% BSA for 1 hour at room temperature. The cells were labeled with Paxillin (5H11) Mouse Monoclonal Antibody (Product # AHO0492) at 2 μg/mL in 0.1% BSA and incubated for 3 hours at room temperature and then labeled with F(ab’)2-Goat anti-Mouse IgG (H+L) Cross-Adsorbed Secondary Antibody, Alexa Fluor® 647 conjugate (Product # A-21237) at a dilution of 1:500 for 1 hour at room temperature. Nuclei were stained with Hoechst 33342 (Thermo, trihydrochloride, trihydrate-10 mg/mL solution in water). F-actin was stained with Phalloidin (Solarbio, Product # CA1610, 1:100). To observe the microstructure of the microcylinder structure, to calibrate the interface, use fluorescent fibronectin. The PDMS replicates were then coated with 50 μM fibronectin in PBS (Cytoskeleton, Inc. Green Fluorescent, HiLyte 488).

### Image acquisition

Here we visualize the cell nuclei and use NLS-GFP for both NIH3T3 fibroblasts and MCF10A epithelial cells. Time-lapse experiments were conducted at 10× and 10× magnification under confocal Leica SP-8 fluorescence microscopy with temperature, humidity, and CO2 regulation (Life Imaging Service) with four-minute interval for 12 hours. Fluorescently marked cells were observed under the same one with a 25× water immersion objective. Due to optical limitations and the finite numerical aperture of the microscope objectives and the limited image quality of the z-axis, the microscopic fluorescent structure on the curvature is imaged by turning the sample upside down.

### Image Analysis

The nuclei of the cells were processed with ImageJ, Imaris, and Matlab routines. Further analysis was occasionally performed on Origin. Convert the position coordinates under the cylindrical coordinates of the curvature side into two-dimensional plane coordinates to calculate the corresponding feature quantity.

### Cell migration Analysis

The time-dependent 3D trajectories of the centroids of the individual cells migrating on inner microcylinders were firstly transformed into 2D coordinates ***r***(*t*) = [*z*(*t*), *R*_*S*_*φ*(*t*)) (see Fig. S2 in the Supplementary Information). The 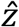 and 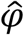 are denoted as the axial and lateral direction of the microcylinders. The cell displacements were calculated by *d****r***(*t*) = ***r***(*t* + *dt*) − ***r***(*t*) at the smallest time lag *dt* = 4min (timeframe interval). The cell migration velocity was calculated using ***v***(*t*) = | *d****r***(*t*)/*dt*|. The cell migration orientation was calculated using *θ*(*t*) = tan^−1^(*v*_1_/*v*_*z*_). As for the cell migrating on planar surfaces, we can calculate the mean square displacement and velocity correlation function to obtain the persistent time *τ*_*p*_. Meanwhile, the persistence time on planar surfaces and quasi-2D (inner microcylinders) was also defined as the time needed for a cell to change its original direction of motion by 90°. All trajectory analyses were performed using a written Jupyter Notebook. Unless otherwise specified, error bars in the figures represent the SEM. Cell position data under all curvature conditions are repeated multiple times.

## Acknowledgements

We thank Samuel Safran for many useful discussions. X.X. is supported by National Natural Science Foundation of China (NSFC, No. 12004082), and by 2020 Li Ka Shing Foundation Cross-Disciplinary Research Grant (No. 2020LKSFG08A). X.Y., Q.F., X.W., and F.Y. are supported by the National Key Research and Development Program of China (2020YFA0908200), by the National Natural Science Foundation of China (NSFC, Grant Nos. 12074407, 12090054, T2221001), by Strategic Priority Research Program of Chinese Academy of Sciences (Grant No. XDB33000000), and by the Youth Innovation Promotion Association of CAS (No. 2021007).

## Author contributions

X.X., Q.F., and X.W. conceived the research. H.W. and X.X. developed the theory. X.Y., Q.F., X.W., and F.Y. designed and carried out the experiments. All authors analyzed the data and wrote the paper. X.Y. and H.W. contributed equally to this work.

## Additional information

The authors declare no competing financial interests. Supplementary information is available in the online version of the paper. Reprints and permissions information is available online at www.nature.com/reprints. Correspondence and requests for materials should be addressed to X.X., or Q.F, or X.W.

